# Microbial lipid shifts in a multi-stage simulated gut

**DOI:** 10.1101/2025.09.25.678496

**Authors:** Ifrat Tamanna, Katja Salonen, Helena Mannochio-Russo, Matilda Kråkström, Paulo Wender P. Gomes, Vincent Charron Lamoureux, Ipsita Mohanty, Sofia D. Forssten, Arthur C. Ouwehand, Tuulia Hyötyläinen, Alex M Dickens, Pieter C Dorrestein, Matej Orešič, Santosh Lamichhane

**Affiliations:** Research Center for Infections and Immunity, Institute of Biomedicine, University of Turku and Turku University Hospital, Turku, Finland; Turku Bioscience, University of Turku and Åbo Akademi University, Turku 20520 Finland; Collaborative Mass Spectrometry Innovation Center, Skaggs School of Pharmacy and Pharmaceutical Sciences, University of California San Diego, La Jolla, CA, USA; Faculty of Chemistry, Federal University of Pará, Belém, PA, Brazil; IFF Health Sciences, 02460 Kantvik Finland; Department of Chemistry, Örebro University, 70281 Örebro Sweden; Faculty of Medicine and Health, Örebro University, 702 81 Örebro Sweden

## Abstract

Food residues that bypass human digestion are further digested by gut microbes, leading to the production of diverse metabolites, including lipids. To investigate how lipids are affected during this transition, we used a colon simulator with four distinct vessels that mimics the proximal to distal part of the human colon. We observed dynamic shifts in a diverse array of microbially derived lipid molecules in the simulated intestinal chyme, including bile acids and *N*-acyl amides with short and odd-chain lipids. Histamine-linked *N*-acyl lipids increased from the proximal to the distal colon vessels (pH 5.5 - 7.0), whereas putrescine-linked, initially abundant in the media, decreased across the colon vessels. We uncovered dynamic associations between in vitro-derived short-chain *N*-acyl lipids and major lipid species such as cholesterol esters, phosphatidylethanolamines, ceramides, and sphingomyelins. To determine the broader relevance of these findings, we applied a reverse metabolomics approach and examined lipid profiles in human small intestine and fecal samples from public datasets. This validated the colon simulator as a model for studying diet-derived and microbially transformed metabolites with relevance to human and animal health and could perhaps be used as a strategy to discover microbial metabolites.

## Introduction

When food enters the stomach, it is transformed into chyme, a biofluid composed of partially digested food and stomach and small intestinal secretions, which then moves into the lower digestive tract. There, gut microbes continue the digestive process, resulting in a plethora of downstream metabolites ^1,2^. The pool of gut microbial-derived compounds and how these microbial metabolites influence host physiology remains underexplored. Water-soluble, polar compounds such as short-chain fatty acids (SCFA), trimethylamine-linked compounds (e.g., trimethylamine *N*-oxide), tryptophan-derived metabolites (e.g., indole, indole-3-acetic acid, indole-3-aldehyde, tryptamine), secondary bile acids (e.g., deoxycholic acid, lithocholic acid), sphingolipids, phenolic and aromatic compounds (e.g., phenylacetic acid, *p*-cresol, 4-hydroxyphenylacetic acid, phenylpropionic acid), and polyamines such as putrescine and cadaverine have been the primary focus of microbe-host health research ^1,3–5^. However, despite the recognized importance of lipids in cellular signaling, structure, and function across both prokaryotes and eukaryotes, microbe-derived lipids remain largely underexplored, except for bile acids and short-chain fatty acids ^6,7^. Emerging evidence indicates that the gut microbiota transforms a wide range of lipids that, in turn, regulate the host lipid homeostasis ^8^. Beyond modifying bile acids, which fall within the lipid molecular class, recent studies have uncovered that gut microbes also influence various other lipid profiles, including the composition of *N*-acyl lipid pools ^9^, cholesterol ^10,11^, sphingolipid ^12,13^, which are gaining attention for their roles in human health. Furthermore, studies also highlight those hundreds of thousands of microbial lipids are yet to be discovered ^8,14^.

Stool metabolome offers valuable insights into the functional roles of gut microbes ^15^. However, our understanding of the biogeographical distribution of these microbial compounds, in particular gut lipids, remains limited. An in vitro gut model provides a controlled experimental condition and dynamic sampling for the characterization of gut microbiota-derived metabolites ^16,17^. In this study, we utilized intestinal chyme from the colon simulator using high-resolution tandem mass spectrometry and characterized the dynamics of emerging, microbially modified lipids within the simulated colon biogeography.

## Results and discussion

Here, metabolites were characterized in the simulated intestinal chyme samples from a colon simulator designed to replicate human colon conditions. The simulator’s four vessels (V1-V4) represent the proximal to the distal part of the large intestine, varying in pH, flow, and anaerobically (Figure 1) ^1,18^. We analyzed 44 simulated chyme samples from these four vessels and the media using an MS/MS-based untargeted metabolomics approach, and metabolites were annotated using the GNPS public spectral libraries, SIRIUS, and an in-house library which consists of masses, retention times, ion mobility values and fragmentation spectra ^19,20^. Here, we also used human feces as a quality control sample. Additionally, we combined the MS/MS feature with reverse metabolomics to further explore the physiological relevance of annotated features in the colon simulator ^8,21^.

**Figure 1.**
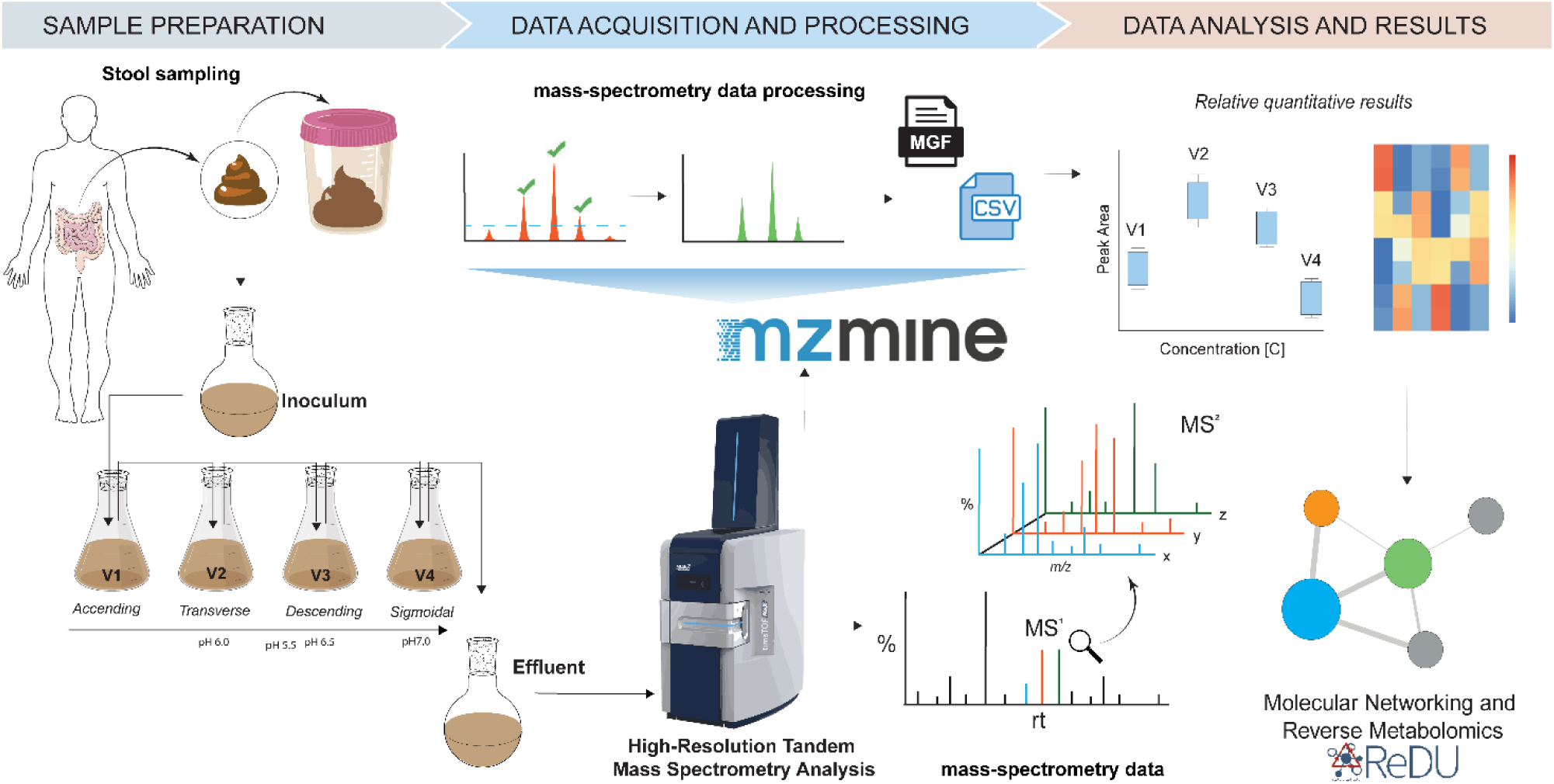
An overview of the sample and the study design. A total of 44 simulated intestinal chymes (11 per vessel) were collected and analysed from the colon simulator. The vessels in one unit (V1–V4) simulate the different compartments of the human colon from the proximal to the distal part, each having a different controlled pH and flow rate. The whole unit is maintained anaerobically and at 37°C. Tandem mass spectrometry-based metabolomics of the simulated intestinal chyme was performed using liquid chromatography-ion mobility spectrometry.

### Untargeted analysis in the simulated intestinal chyme

To explore the dynamics of lipid-linked molecules in the simulated fecal samples, we analyzed the untargeted MS/MS data obtained from these simulated fecal samples using Feature based molecular networking in GNPS and SIRIUS. Of the 6,903 MS/MS features obtained after processing in MZmine, we were able to annotate 750 features within the GNPS libraries (≈10.86%, Figure 2a, annotated features are highlighted in green), of which 128 features were related to the candidate feature annotations ^14^ and 461 were suspect-related matches. Sirius-based CANOPUS ^22^ predicted 17 compound classes from 5,383 (≈96.38%) MS/MS features (Fig 2b), where 1,865 features were linked with lipids and lipid-like molecules and annotated features ^23^. Next, we sought to analyze the dynamics of these annotated features in different parts of the colon simulator with reference to media. Of the analyzed annotated and lipid-like molecules, 1,025 (391 GNPS annotated, Figure 2c and S1, Supplementary Table 1 and 2) features showed a significant change in at least one of the vessels and/or in human stool samples when compared to the media.

**Figure 2.**
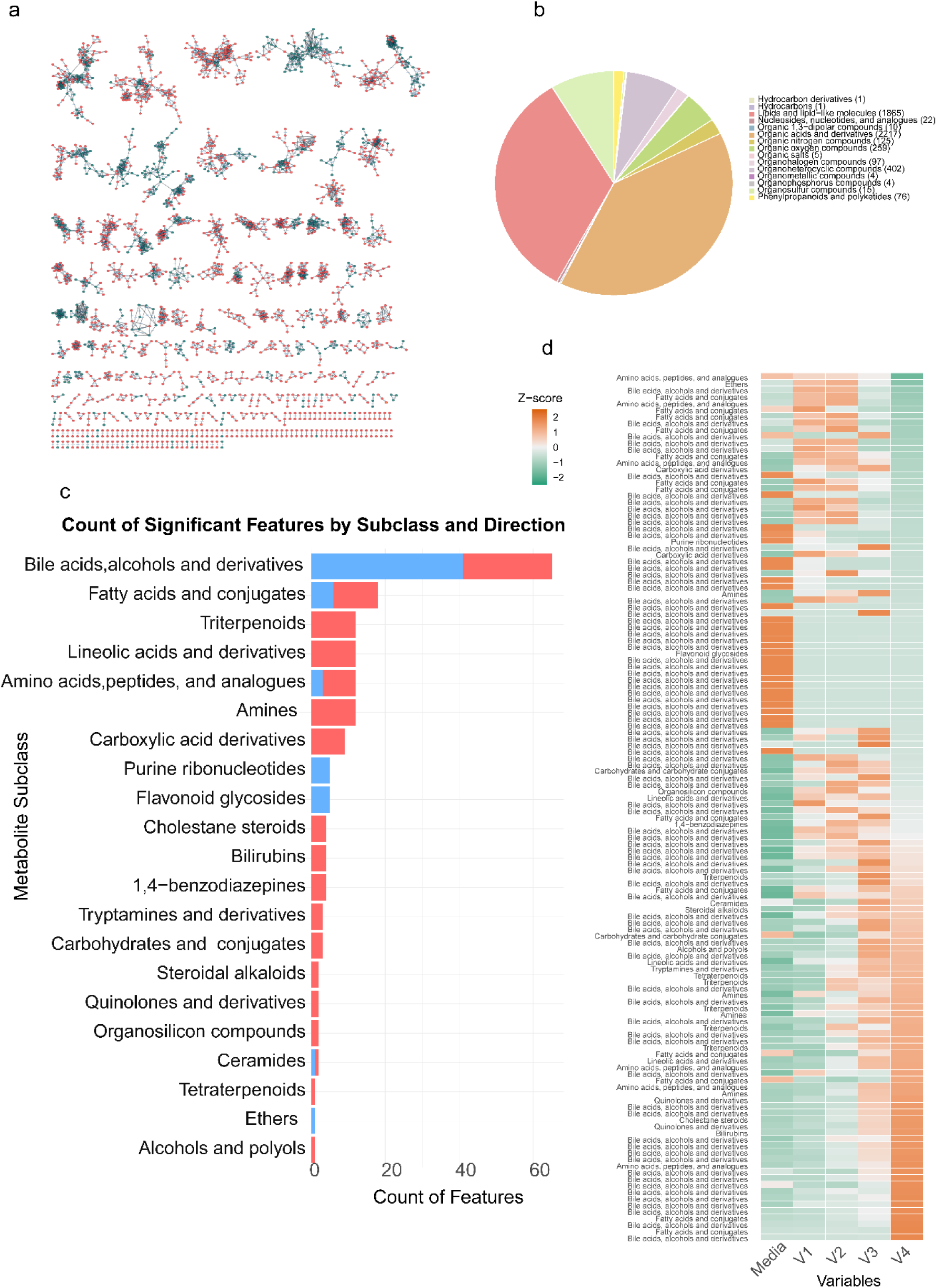
Dynamics of lipids and lipid-like molecules in the gut simulator. a) In the network, we show the dynamics portion of both known and unknown compounds relating to colon simulators, which have a potential link with microbial metabolism. Green in the network indicates annotations from GNPS, while red is unknown. b) Compound class predicted from Sirius. c) Bar plot showing the count of significant annotated metabolic features within each metabolite subclass, separated by effect direction based on the sign of the coefficient from the metric -log(qval) * sign(coef), which reflects both statistical significance and effect direction. Subclasses are ordered from top to bottom by the total number of significant features, highlighting the most impacted metabolite categories. Red bars indicate metabolites class with positive effects, and blue bars indicate metabolites with negative effects, as determined by the sign of the coefficient and significance of the q-value. d) Heat map showing the annotated metabolic features within each metabolite subclass. This plot highlights bile acids and fatty acid derivatives are the most impacted metabolite categories included based on sirius. Variables were reordered based on descending z-scores in V4, with ties further sorted by descending z-scores in Media

The majority of these changes include the bile acid subclass and the fatty acid and derivatives, including *N*-acyl lipids, features of odd fatty acid chains linked to lipids Leucine-C19:0, Leucine-C21:0, Leucine-C21:1, potentially of microbial origin (Figure 2c and Supplementary Figure S1). Here, bile acid features are matched to the recently described bile acid amidates, including Gln-HDCA, Glu-gMCA, dihydroxylated, candidate tri-as well as tetrahydroxylated bile acid, which are reported to be not usually at detectable levels in healthy adult humans (except infants). Post-molecular networking MassQL query validated three tetrahydroxylated bile acid features (https://massqlpostmn.gnps2.org/, Supplementary Table 3, Supplementary Figure 2c). Two out of three of these bile acids showed an increased pattern from the proximal vessel to the distal part of the colon. Gut microbes are established regulators of bile acid metabolism, particularly in transforming primary bile acids into secondary forms through deconjugation and dihydroxylation. Tetrahydroxylated bile acids are a distinct class of bile acids characterized by high hydrophilicity and minimal cytotoxicity. We acknowledge that the observed candidate tetrahydroxylated bile acids annotation is based on MS/MS feature match, which needs to be further validated. However, our results, along with reverse metabolomics-based MASST analysis, suggest a plausible connection to the gut microbiota and polyhydroxylated bile acid (Figure S2). Other annotated features from diverse metabolite classes, such as fatty acid and novel conjugates of lipids, amines, amino acids, steroids, terpenoids, flavonoids, and various small organic compounds that were significantly altered (p < 0.05) in at least one vessel are shown in Figure 2 c-d, Supplementary Table 1-2.

### *N*-acyl amides in the colon simulator

We observed that the MS-MS features of fatty acids and their derivatives, including newly discovered *N*-acyl lipids, were altered in different parts of the colon simulator. Next, we focused on investigating potential microbial *N*-acyl lipids, which are fatty acids linked to amine groups via amide bonds ^9^. Using an in-house library, we identified 52 *N*-acyl lipids in the simulated chyme slurry samples and human inoculum. Among these, the most prominent were amino groups conjugated with long carbon chains (greater than 12 carbons), such as C18:0, followed by C18:1 (Figure 3a). Additionally, a small number of amino groups were linked to short carbon chains (fewer than 6 carbons in length). Notably, Lys-C17:0 emerged as the most abundant *N*-acyl amide across all samples, followed by GABA-C20:0. We next evaluated the variability of *N*-acyl amides across different vessels compared to the media. To understand the dynamics and distribution of *N*-acyl amides in each vessel, we calculated the mean value of each *N*-acyl lipid variable within each vessel and media group. Then, we compute the z-score of these means across vessels and media for each variable as shown in Figure 3b. We found a clear pattern among the conjugated lipids, with the most prominent trend of these *N*-acyl amides showing increased levels towards the distal part of the colon (Figure 3b), suggesting conjugation or degradation pathways may be more active in some part of the intestine than others. In contrast, conjugates with putrescine headgroup had higher abundance in the media, which remained low in vessel 1 and relatively stable throughout the simulation. A multivariable linear model analysis revealed that four *N*-acyl amides were significantly altered (p < 0.05) in at least one vessel, with a false discovery rate threshold of < 0.25 (Supplementary Table 1). Specifically, levels of histamine-C5:0 and Arg-C18:0 increased from vessel 1 to vessel 4 over the course of the simulation (Figure 3, Supplementary Figure S4). Histamine controls gastric acid secretion in hosts ^24^ and is a monoamine synthesized from the amino acid His mediated via enzymes known as amino acid decarboxylases, enzyme also found in members of the gut microbiome ^25^. Intriguingly, we also found that the level of His-C8:0 conjugate was lower in media and proximal vessels 1 and 2, which gradually increased in the subsequent distal vessels (Figure 3e), suggesting ongoing microbial activity. In contrast, putrescine-C19:2 remained high in media, low in vessel 1, and relatively stable throughout the simulation (Figure 3d) while ornithine and Arg conjugates showed the reverse trend. It is noteworthy that Arg and ornithine are the main precursors for putrescine synthesis ^26^, and that gut microbiota play a major role in determining polyamine levels in the lower part of the intestine ^27^.

**Figure 3.**
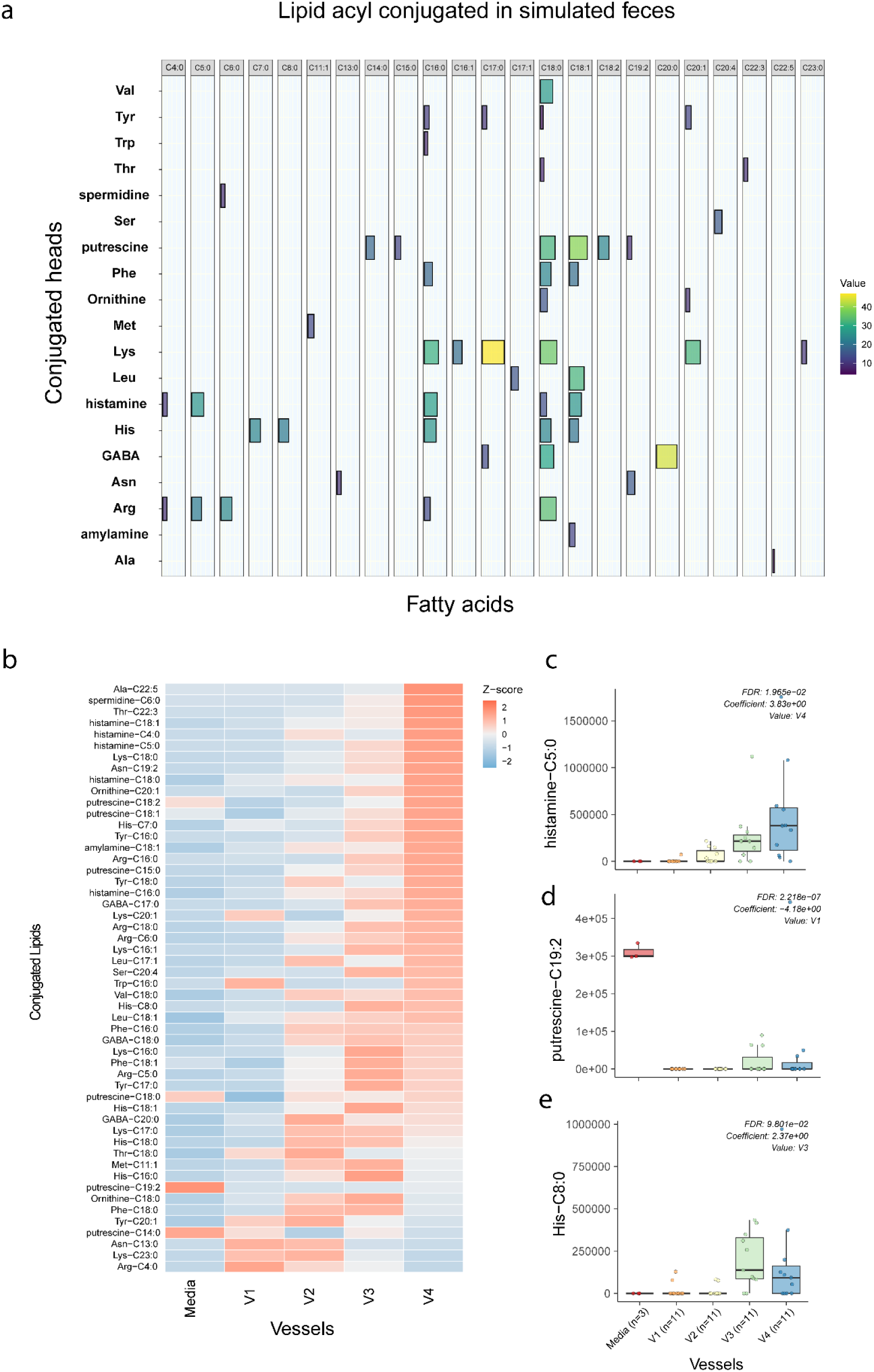
*N*-acyl amides detected in the colon simulator. a) A total of 52 conjugated *N*-acyl lipids were detected in colon simulation samples. b) Dynamics of these conjugated *N*-acyl lipids compared with media and vessels. We calculated the mean value of each *N*-acyl lipids (variable) within each vessel and media group. Then, we compute the z-score (standardized value) of these means across vessels and media for each variable. c-e) Box plot showing statistically significant associations of the compared *N*-acyl lipids in at least one vessel or media.

Next, we compared the impact of simulation time 24 hours (n=12) or 48 hours (n=32) and carbon source with (n=12) or without (n=32) PDX (polydextrose, a synthetic dietary fiber). To assess this, we performed multivariate linear modeling (*N*-acyl lipids ∼ Time + Treatment). We found that simulation time had a significant impact on the *N*-acyl lipid profile (Figure 4, Supplementary Table 4), whereas treatment (with or without PDX) did not show a statistically significant effect. We observed that short-chain *N*-acyl lipids (C4-C6) increased with time. We found that histamine conjugates increased, while His conjugates decreased with simulation time, further supporting our previous observation in relation to vessels and microbial activity. We also observed that odd chain conjugate Met-C11:1, and long chain C16 and C18 conjugated to Lys and Phe decreased with increasing simulation time. We found that the synthetic dietary fiber PDX did not have a statistically significant effect, with the exception of putrescine conjugates; however, we observed a lowering trend in short-chain histamine conjugates and an increasing trend for most of the long-chain conjugates. We acknowledge limitations in our study, particularly the small and uneven sample sizes for the time and treatment groups. Nonetheless, our study uniquely reports the dynamics of these emerging *N*-acyl lipids in different sections of the colon. Our study corroborates the potential role of microbes in the conjugation of these lipids.

**Figure 4.**
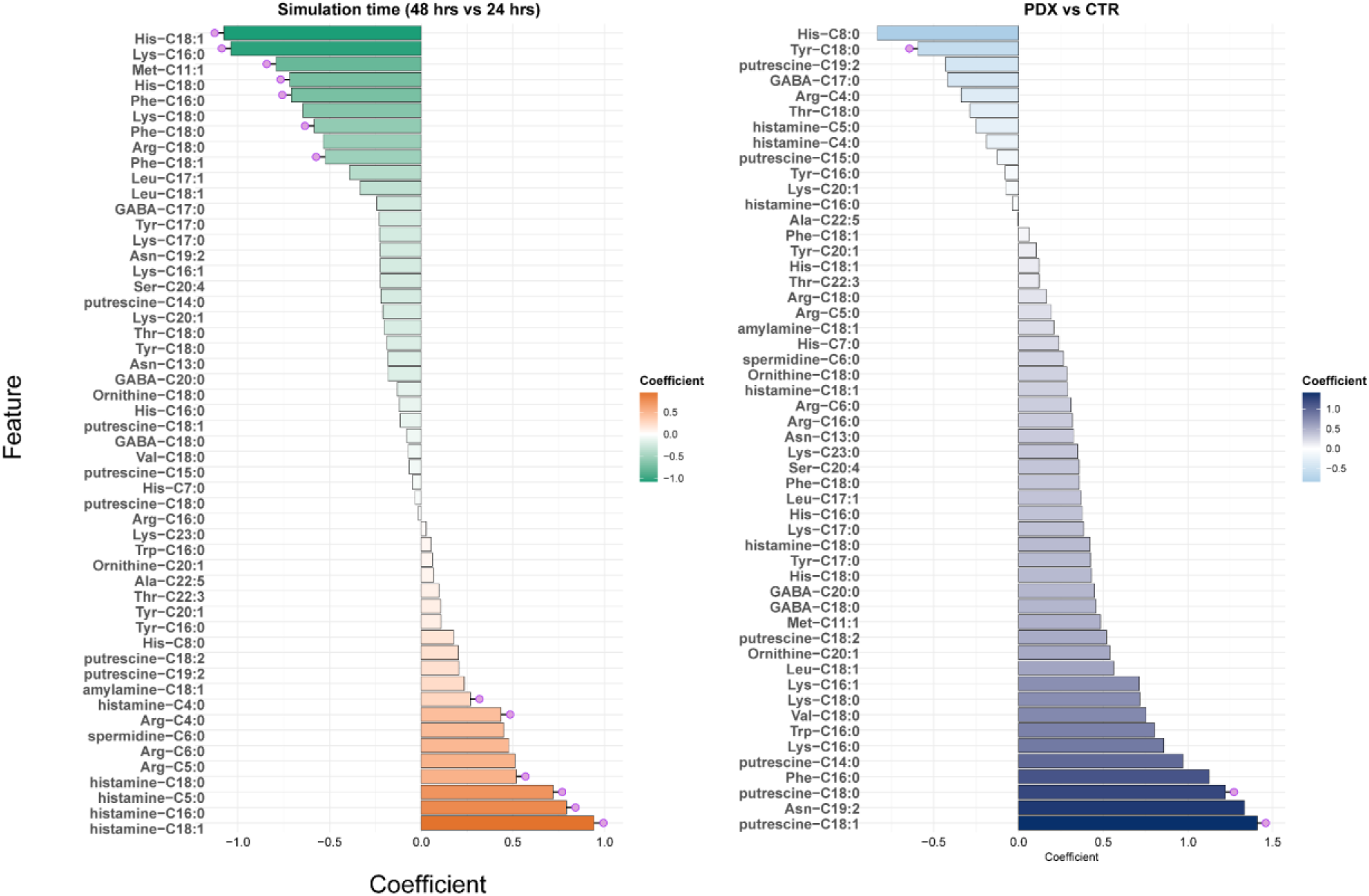
Impact of the simulation time (24 vs. 48 h) and PDX treatment on the *N*-acyl lipid profile. Forest plot illustrating the coefficient estimate of a linear model (*N*-acyl lipids ∼ Time + Treatment). Red and deep blue bars represent positive correlations, while green and muted blue bars represent negative correlations, as determined by linear regression models. The p-values shown are nominal; adjusted p-values (corrected for multiple comparisons using the Benjamini-Hochberg method) are available in Supplementary Table 5.

The latest study also shows that the gut microbiome diversifies *N*-acyl lipid pools, including those derived from short-chain fatty acids, especially linked with histamine and polyamine conjugates, which are linked to HIV status and cognitive impairment ^9^. In particular, the short-chain histamine conjugates were produced by microbial monocultures of *Collinsella aerofaciens* ATCC 25986, *Holdemanella biformis* DSM 3989, and *Prevotella buccae* D17. Intriguingly, there are reports on the functions and therapeutic potential of *N*-acyl amino acids ^28^, which are endocannabinoid-like molecules. The connection between the endocannabinoid and the gut has been recognized for over half a century ^29^. As early as the 1970s, studies demonstrated that endocannabinoids can significantly influence gut motility ^29^. More recent research suggests that gastrointestinal transit time can, in turn, shape the composition and function of gut microbiota. In our study, *N*-acyl lipid compounds similar to endocannabinoids differed between the beginning and end of the colon simulator. This suggests there may be a temporal and spatial connection between endocannabinoid activity and the gut microbiome.

To further understand if these same metabolites are found in animal and human samples, these statistically altered *N*-acyl lipids in different vessels, histamine-C5:0, Arg-C18:0, putrescine-C19:2, and His-C8:0 were subjected to reverse metabolomics ^30^. In reverse metabolomics, MS/MS spectra, in this case for the abovementioned *N*-acyl lipids, are searched across thousands of studies, and the metadata associated with files that have matching MS/MS spectra can be used for biological interpretation ^8^ ^21^. The data sources we searched include MetaboLights, Metabolomics Workbench, and GNPS/MassIVE ^31^. We mainly found hits for histamine-C5:0 and Arg-C18:0. Figure 5a shows that these compounds were indeed mainly found in the gut of both rodents and humans. In rodents, these compounds were present in the cecum, jejunum, duodenum, and ileum, while in humans, they were observed in the oral cavity and colon. In addition to the intestine, these compounds were also detected in the pancreas and spleen of rodents. Besides body parts, we also found histamine-C5:0 and Arg-C18:0 was associated with host disease data in the public repositories. Recent work also suggests that short-chain *N*-acyl lipids modulate T cell differentiation ^9^, which in turn can affect overall immune homeostasis. It remains to be seen if these molecules exhibit such function in IBD.

**Figure 5.**
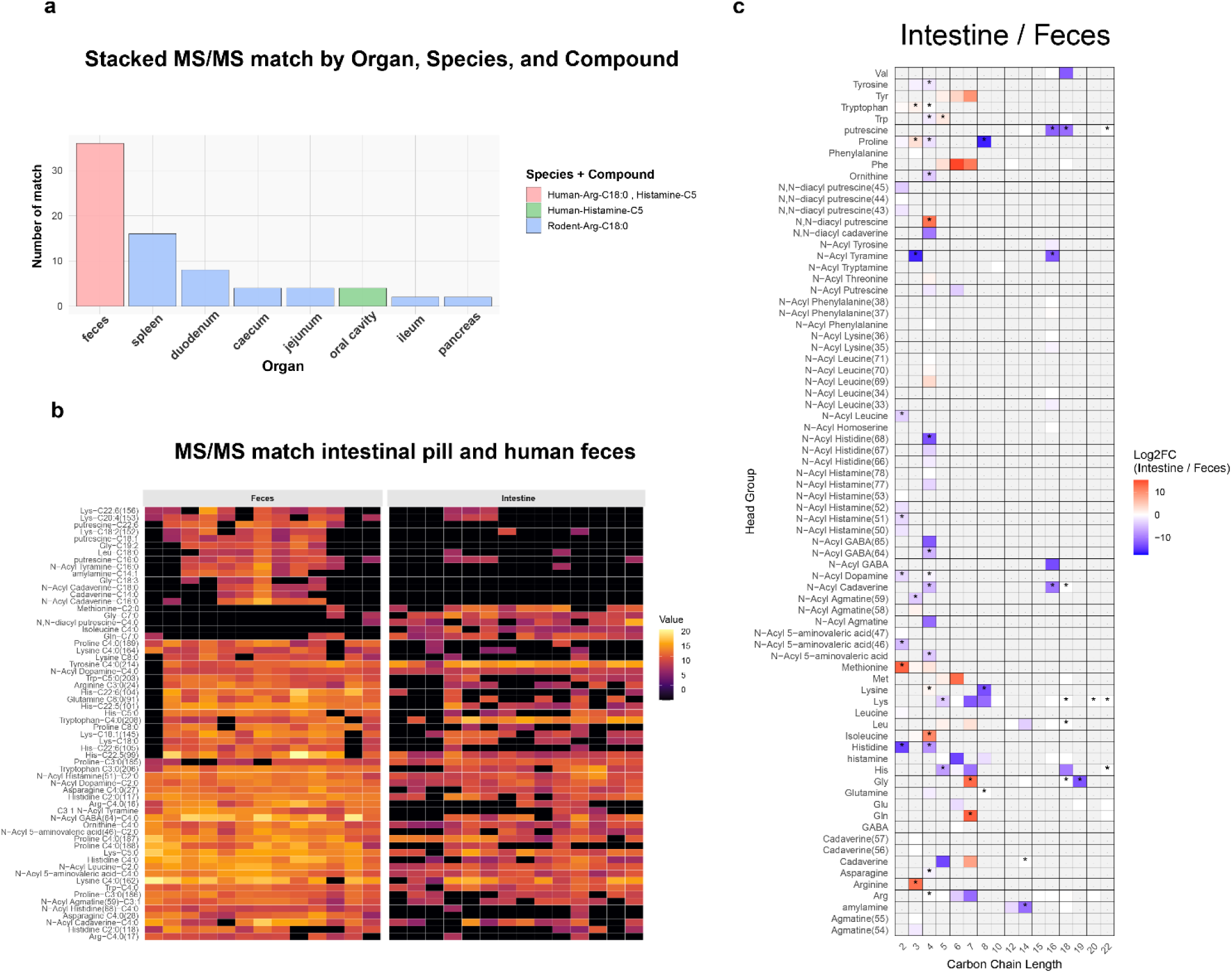
a) Reverse metabolomics of the MS/MS spectral matches for histamine-C5:0, Arg-C18:0, putrescine-C19:2, and His-C8:0. b and c) Distribution of N-acyl lipids between small intestinal fluid and fecal samples. Longer-chain N-acyl lipids such as putrescine–C18:1, Lys–C22:6, Gly–C19:2, and Leu–C18:0 were enriched in fecal samples, whereas small intestinal fluid contained higher levels of methionine–C2:0, isoleucine–C4:0, and N, N-diacyl putrescine–C4:0. Statistical tests were performed using the non-parametric Mann-Whitney U test when two groups (feces vs. intestine) were compared. The p-values were corrected for multiple comparisons using the Benjamini-Hochberg correction. In the panel (c), Fold change for the 217 MS/MS spectra matched to N-acyl lipids was calculated as the ratio of median peak areas in Intestin versus Feces samples with a pseudocount of 1 added to avoid division by zero. FC = (median_Intestin + 1) / (median_Feces + 1). Here, grey indicates that data is not available/missing.

To further contextualize the biogeographical relevance of N-acyl lipids *in vivo* in humans, we re-analyzed a publicly available untargeted LC–MS/MS dataset (MSV000094551) comprising paired small intestinal fluid and fecal samples from healthy participants ^32,33^. Figure 5 b and c and Supplementary Figure S5 illustrate the distribution of N-acyl lipids across these two sample types. MS/MS spectra were matched against the GNPS spectral libraries, including the recent MassQL-derived N-acyl lipids ^34^ and candidate bile acid libraries^14^. In total, 217 MS/MS spectra matched to N-acyl lipids in this dataset (Supplementary Figure S5, GNPS taskID: 6ab06bfcdd27479199c1c7d5231d8227). Peak intensity analysis with available metadata revealed that 62 of these matched N-acyl lipid spectra significantly differed between small intestinal fluid and fecal samples (Figure 5b), consistent with in-vitro observation of ongoing microbial activity along the gastrointestinal tract. Here, in particular longer-chain N-acyl lipids such as putrescine–C18:1, Lys–C22:6, Gly–C19:2, and Leu–C18:0 were enriched in fecal samples, whereas small intestinal fluid contained higher levels of methionine–C2:0, isoleucine–C4:0, and N, N-diacyl putrescine–C4:0.

### Microbial-derived conjugated *N*-acyl amides associate with major lipid classes

We also further sought to understand how microbial-derived conjugated *N*-acyl amides interact with major lipid classes and endocannabinoids. For this analysis, we performed a total lipid class-specific association study with the *N*-acyl amides in the colon simulator (Figure 6). In this approach, individual lipid concentrations within each lipid class were summed, and subsequent data analysis was conducted by treating each lipid class as a single variable. Overall, we found a positive association pattern between most lipid classes and conjugated *N*-acyl amides from vessel 1 to vessel 4, with the exception of lysophosphatidylethanolamine, which showed a more negative pattern toward distal vessel (V4). Notably, short-chain *N*-acyl amides (fewer than 8 carbons in length), such as C4:0, C5, C6, and C7 (SCFA conjugates), exhibited a strong dynamic association from vessel V1 to V4, particularly with cholesterol esters, phosphatidylethanolamines, ceramides, and sphingomyelins. In contrast, the association pattern for long-chain conjugates (more than 16 carbons in length) was less pronounced across the vessels, except for phosphatidylglycerol and lysophosphatidylethanolamine, indicating gut microbial activity driven to SCFA-diversification, suggesting certain fatty acids being conjugated more readily. Similarly, for endocannabinoids, we observed a pattern resembling that of the summed lipid classes. Conjugates with short carbon chains were positively associated with overall *N*-acyl amide levels detected in the simulated chyme, particularly oleoylethanolamide, stearoylethanolamide, and palmitoylethanolamide in vessels V3 and V4. This trend was less pronounced for arachidonic acid and 2-arachidonoylglycerol (2-AG). Notably, associations were strongest in vessel V4, except for 2-arachidonoyl glycerol ether, which showed a negative association, including with long-chain *N*-acyl amide conjugates. In contrast, 2-AG exhibited a clear positive association with most short-chain *N*-acyl amide conjugates from V2 to V4, including histamine-C5 (Supplementary Figure S6), which varied significantly across the different vessel sections in the colon simulator. Changes in these association patterns across vessels imply differential metabolism, uptake, or biotransformation of lipids within the simulator.

**Figure 6.**
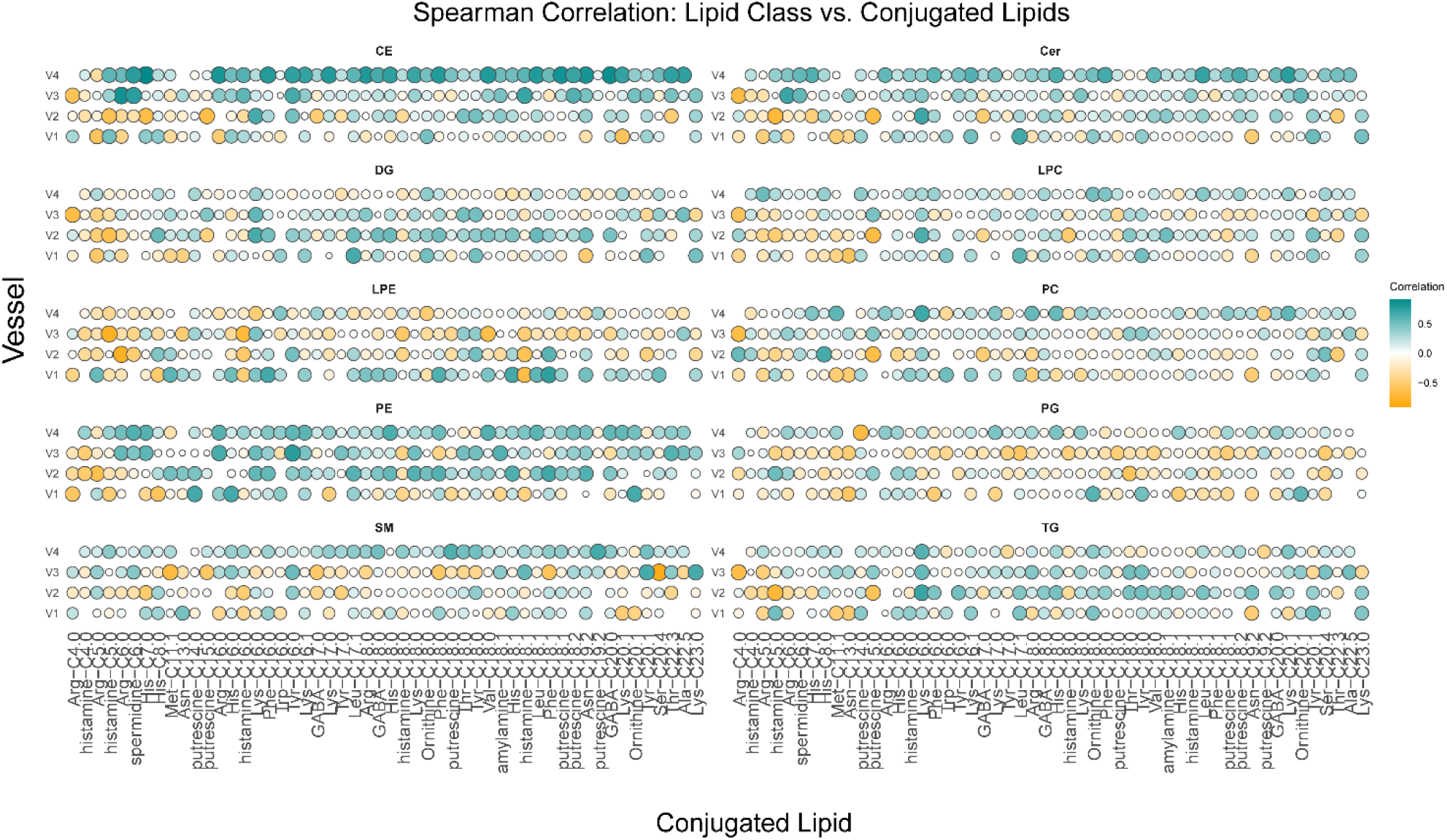
Association of the novel lipids with the lipidome in the colon simulator. Spearman correlation coefficients illustrated by heat map. Lipids class-wise correlated with the *N*-acyl lipids in the simulator’s four vessels (V1-V4) that represent the proximal to the distal part of the colon. The lipid classes: triacylglycerols (TG), ceramides (Cer), cholesterol esters (CE), diacylglycerols (DG), lysophosphatidylcholines (LPC), phosphatidylcholines (PC), phosphatidylethanolamines (PE), and sphingomyelins (SM).

## Conclusions

This study highlights the dynamics of non-polar microbial lipids, including newly discovered microbial *N*-acyl lipids, across distinct simulated gut regions, emphasizing an underexplored class of metabolites in microbiome-focused metabolomics. This study shows that a colon simulator can effectively model the dynamic changes in microbially derived lipids along the gut, revealing region-specific shifts from the proximal to distal colon. Notable transitions were observed in bile acids, *N*-acyl amides in relation to classical lipids. The major limitation of this study is the absence of microbiome data, which prevents us from establishing a direct causal link between gut microbiota and lipid profiles. Nevertheless, our lipidomics approach combined with the reverse metabolomics-based approach using simulated intestinal chyme serves as a valuable tool for hypothesis generation, enabling the identification of candidate microbial lipids for future investigation. Our findings also highlight the value of a colon model, which offers controlled and dynamic sampling conditions for investigating gut microbiota-associated potential bioactive lipids. Further, exploration through public datasets confirms the in vitro gut model relevance for studying diet and microbe-derived metabolites important to human and animal health.

## Supplementary Figures

**Figure S1.**
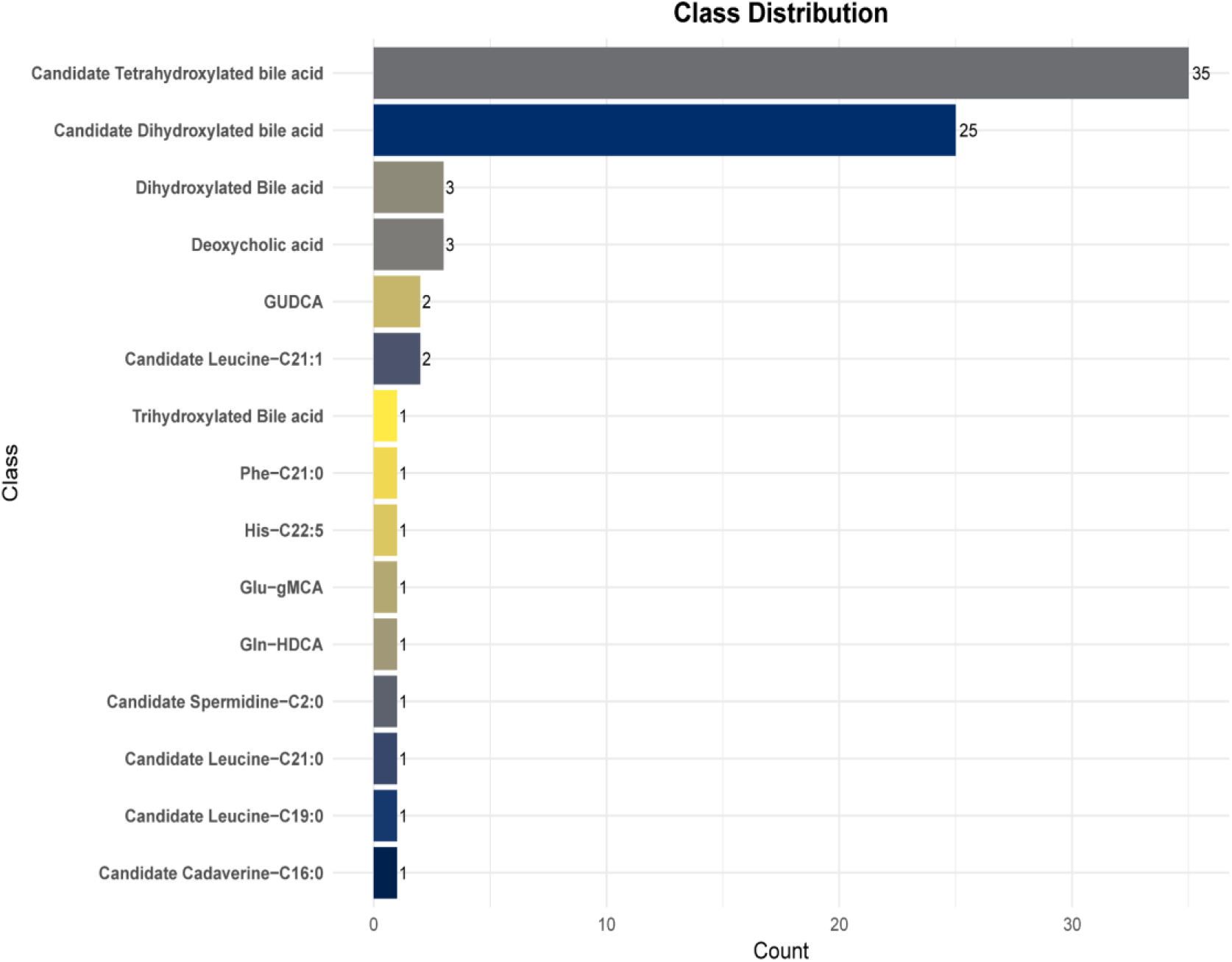
Number of candidate bile acid and the N acyl lipids features that showed change in at least one of the vessels and or in human fecal QC sample in reference to the media. We acknowledge that observed candidate bile acids and other annotations are based on MS-MS feature match, which needs to be further validated.

**Figure S2.**
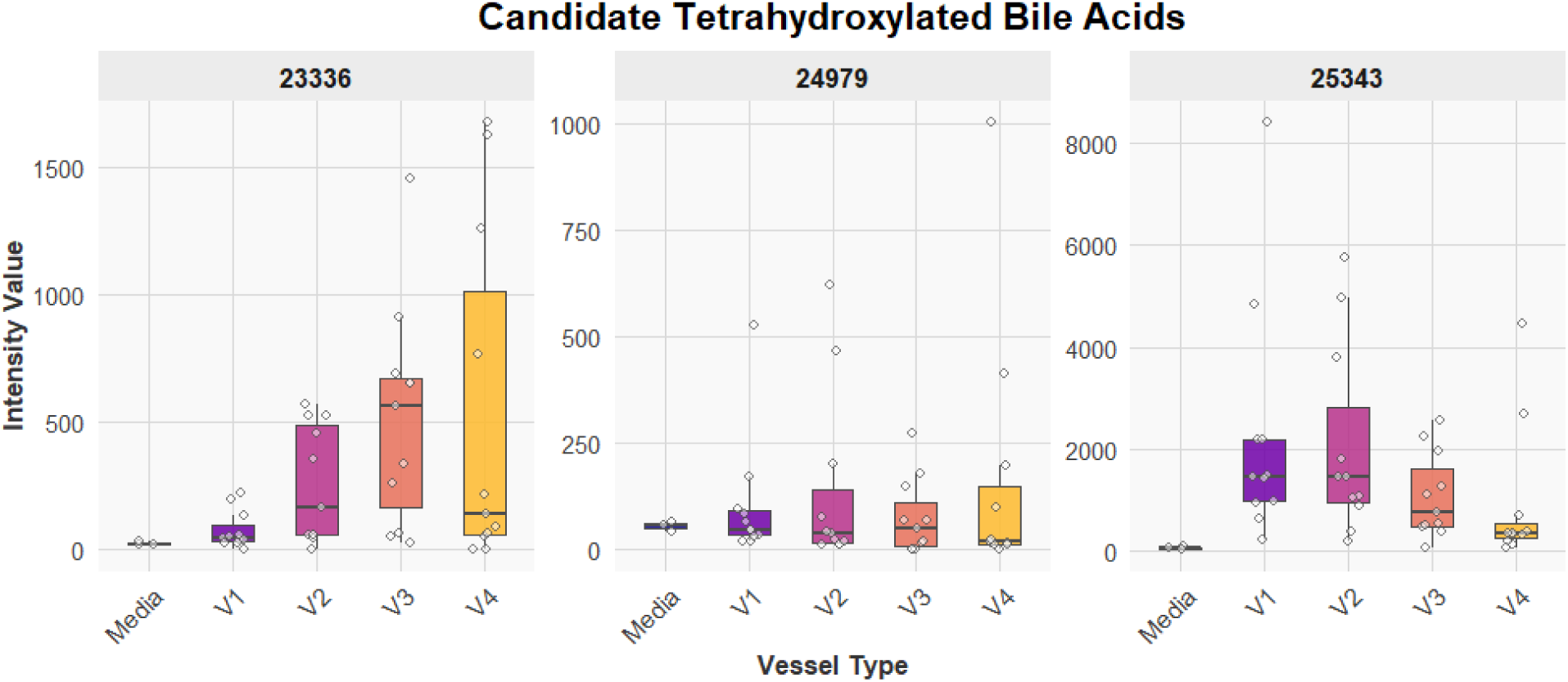
Dynamics of candidate tetrahydroxylated bile acid, the simulator indicates ongoing microbial activity. These selected compounds highlight the trend between the vessels (V1–V4), which mimics the different compartments of the human colon from the proximal to the distal part.

**Figure S3.**
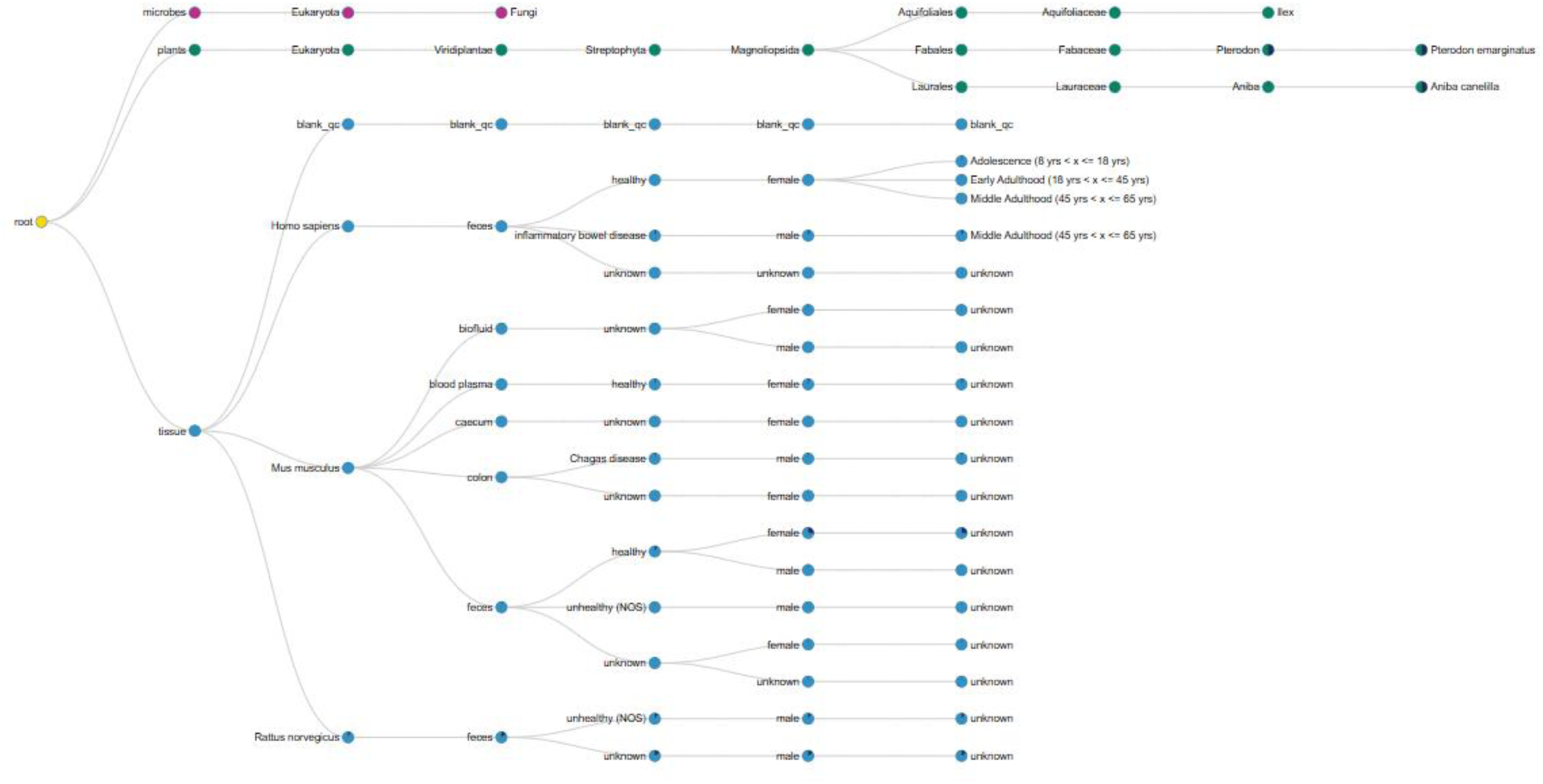
The output of the domain-specific MASST search for tetrahydroxylated bile Facid MS/MS spectra against a reference database of MS/MS data.

**Figure S4.**
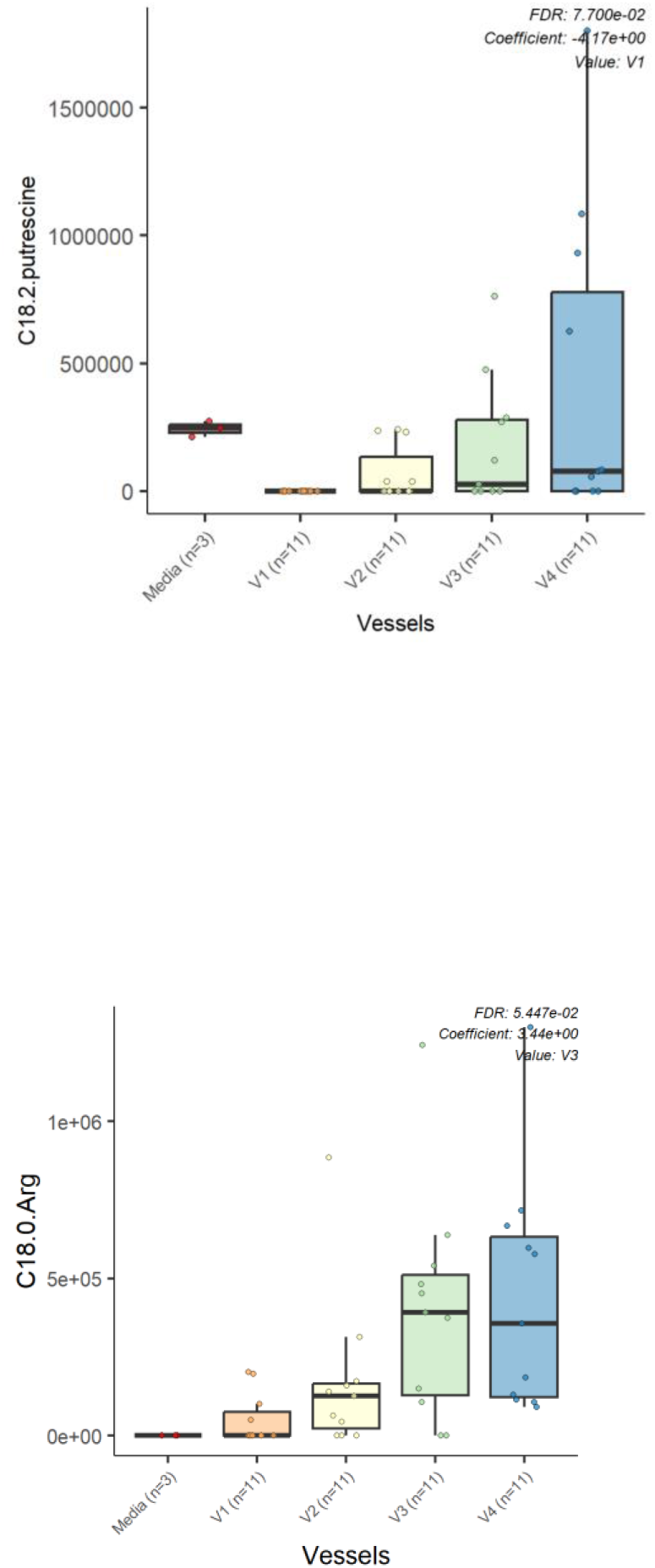
Box plot showing statistically significant associations of the compared *N*-acyl lipids in at least one vessel or media.

**Figure S5.**
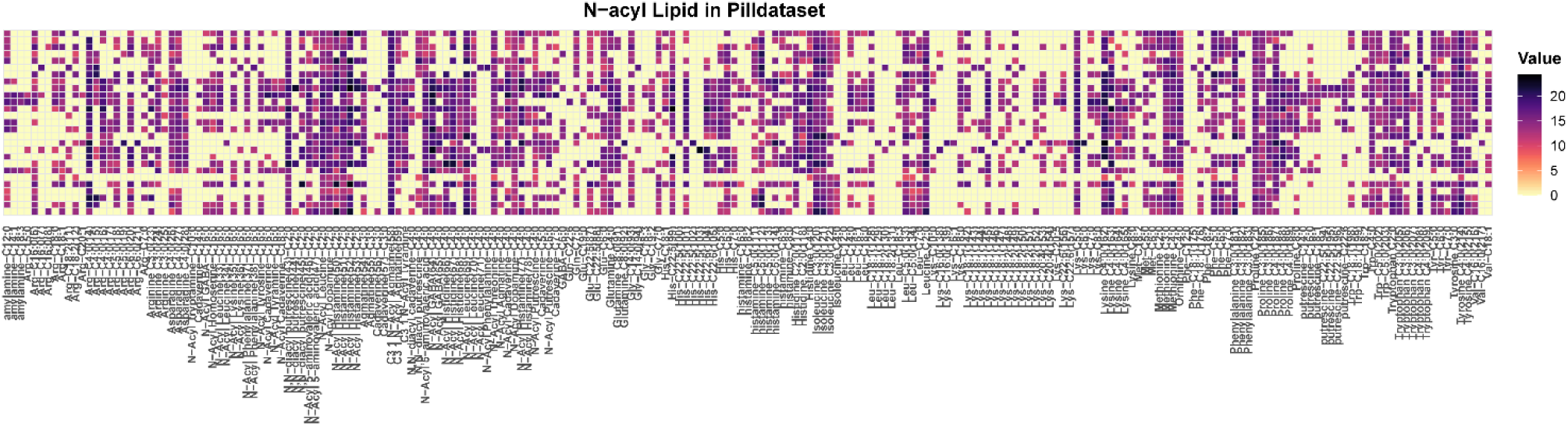
Distribution of N-acyl lipids between small intestinal fluid and fecal samples in a publicly available untargeted LC–MS/MS dataset (MSV000094551)

**Figure S5.**
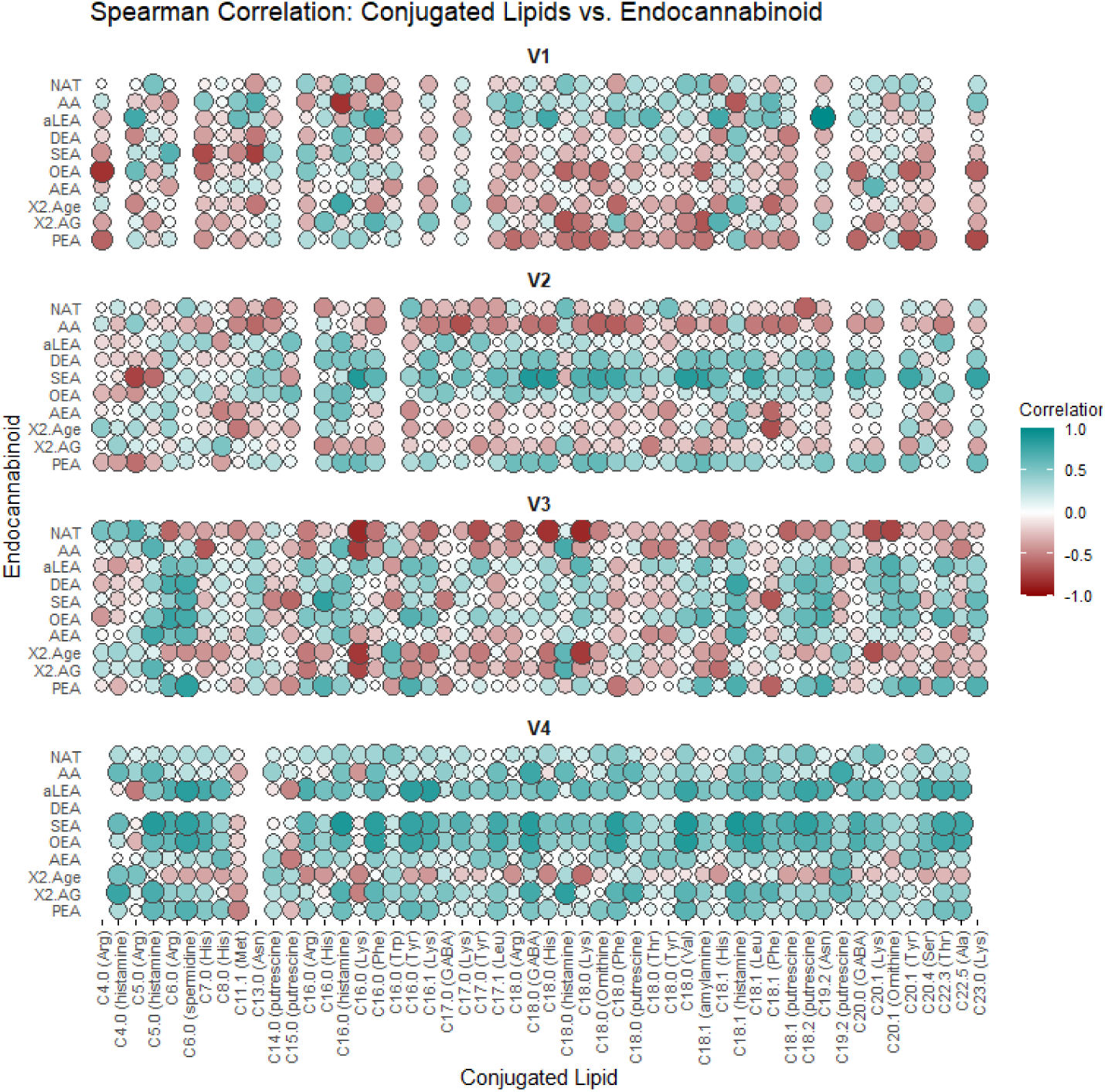
Association with endocannabinoids (ECC) in four vessels. These included palmitoylethanolamide (PEA), arachidonoyl glycerol (AG), 2-arachidonic glycerol ether (2-Age), arachidonoylethanolamide (AEA), oleoylethanolamide (OEA), stearoylethanolamide (SEA), docosatetraenoylethanolamide (DEA), alpha-linolenoylethanolamide (aLEA), arachidonic acid (AA), N-arachidonoyl taurine (NAT).

## Materials and Methods

### Colon simulator

The Enteromix model of the human large intestine was described in detail previously (*ADD REF, example above in comment).* In brief, this simulator consists of eight separate units, each containing four (V1–V4) semi-continuously connected glass vessels. The vessels in one unit mimic the different compartments of the human colon from the proximal to the distal part, each having a different controlled pH and flow rate. Each unit is kept anaerobically and at 37°C. In the initial phase of the simulation, each unit is inoculated with pre-incubated fecal microbes from a fresh fecal sample, which form the microbiota of the colonic model. In the present study, the fecal samples for inoculum were provided voluntarily by three healthy Finnish volunteers. The study and all methods used in it were carried out in accordance with relevant guidelines and regulations, and informed consent was orally obtained from all research subjects. This simulation was performed at IFF Health Sciences, Kantvik, Finland. To understand the lipidomic changes over time, the microbial slurry was collected from all vessels (V1–V4) after 24 and 48 hours with/without PDX supplementation ^17^. In total 44 samples were collected from the colon simulator vessels (V1–V4) and stored at −80 °C prior to lipidomic analysis. In addition, media and inoculum used for the simulation were collected as quality control samples.

### Untargeted metabolomics

Simulated intestinal chyme extracts were prepared by mixing crash solvent, consisting of 0.1% formic acid (FA) in acetonitrile (400 µL), with 200 µL of slurry in a glass vial and briefly vortexed. The mixture was then left to settle at −20 °C for 30 minutes, followed by filtration through a protein precipitation filter plate. The filtrates were collected into 96-well plates with glass inserts. Next, the samples were transferred into glass vials, dried under a nitrogen stream at 35 °C, and resuspended in 50 µL of the final solution (60% water, 20% ACN, and 20% isopropanol). The prepared samples were stored at −80 °C and, after thawing, were briefly vortexed prior to LC-IMS analysis. The instrumentation for liquid chromatography was a Shimadzu Nexera system (Shimadzu, Kyoto, Japan) equipped with a binary pump LC-40D x3, autosampler SIL-40, degasser units (DGU-405 and DGU-403) and column oven CTO-40C. The separation was performed on a Phenomenex Kinetex 1.7 μm, 100 x 2.1 mm XB-C18 100 Å column, with a Phenomenex SecurityGuard ULTRA Cartridge pre-column, held at 40 °C. The injection volume was 10 µL and the autosampler was set to 15 °C. The mobile phase flow rate was 0.4 mL/min and the eluents were 0.1 % formic acid (FA) in water as a phase A and methanol: acetonitrile (5:1, v/v) as a phase B. The gradient started with 70% of phase B and was increased to 85% in 4.5 minutes, followed by a 1-minute increase to 100% and was held for 2 minutes. After each injection, the needle was washed with 10 % dichloromethane in MeOH and isopropanol for 8.6 seconds each. The detection was conducted on a Bruker timsTOF fleX (Bruker Daltonik, Bremen, Germany) equipped with an electrospray ion source and a TIMS device. The instrument was operated in positive mode and the source settings were as follows: nitrogen drying gas with a flow of 10 L/min at 220 °C, nebulizer gas 2.2 Bar, capillary 4500 V and end plate offset was 500 V. The acquisition scheme employed a data-dependent acquisition-parallel accumulation-serial fragmentation mode. Detection was carried out using combinatorially synthesized reference mixtures with qualitative composition. Bruker Metaboscape software was used for data preprocessing.

For lipidomics prepared using a method based on the Folch procedure [15] as detailed by Lamichhane et al. [16]. An internal standard mixture containing 2.5 µg/mL mL 1,2-diheptadecanoyl-sn-glycero-3-phosphoethanolamine (PE(17:0/17:0)), N-heptadecanoyl-D-erythro-sphingosylphosphorylcholine (SM(d18:1/17:0)), N-heptadecanoyl-D-erythro-sphingosine (Cer(d18:1/17:0)), 1,2-diheptadecanoyl-sn-glycero-3-phosphocholine (PC(17:0/17:0)), 1-heptadecanoyl-2-hydroxy-sn-glycero-3-phosphocholine (LPC(17:0)), 1-palmitoyl-d31-2-oleoyl-sn-glycero-3-phosphocholine (PC(16:0/d31/18:1)) and 1,2,3-triheptadecanoyl-sn-glycerol (TG(17:0/17:0/17:0)) was prepared in CHCl_3_:MeOH (2:1, v/v). Six point calibration curves with concentrations between 100 and 2500 ppb in CHCl_3_:MeOH (2:1, v/v) were prepared for 1-Hexadecanoyl-2-octadecanoyl-sn-glycero-3-oethanolamine (PE(16:0/18:1)), octadecenoyl-sn-glycero-3-phosphocholine (LPC(18:1)), Cholesteryl hexadecanoate (CE(16:0)), 1,2-Distearoyl-sn-glycero-3-phosphoethanolamine (PE(18:0/18:0)), N-stearoyl-D-erythro-sphingosylphosphorylcholine (SM(18:0/18:1)) and Cholesteryl linoleic acid (CE(18:2)). The samples were prepared by spiking 10 µL of sample with 10 µL of 0.9% NaCl and 120 µL of internal standard solution. The samples were vortexed and were left to stand on ice for 30 min. Samples were centrifuged (9400 × g, 5 min, 4 °C) and 60 µL from the lower layer was diluted with 60 µL of CHCl_3_:MeOH (2:1, v/v). For the LC separation, a Bruker Elute UHPLC system (Druker Daltonik, Bremen, Germany) equipped with an auto sampler cooled to 10 °C, a column compartment heated to 50 °C and a binary pump was used. A Waters ACQUITY BEH C18 column (2.1 mm × 100 mm, 1.7 µm) was used for chromatographic separation. The flow rate was 0.4 mL/min and the injection volume was 1 µL. The needle was washed with 10% DCM in MeOH and ACN: MeOH: IPA: H_2_O (1:1:1:1, v/v/v/v) + 0.1% HCOOH after each injection for 7.5 s each. The eluents were H_2_O + 1% NH_4_Ac (1M) + 0.1% HCOOH (A) and ACN: IPA (1:1, v/v) + 1% NH_4_Ac + 0.1% HCOOH (B) The gradient is as follows: from 0 to 2 min 35-80% B, from 2 to 7 min 80-100% B and from 7 to 14 min 100% B. Each run was followed by a 7 min re-equilibration period under initial conditions (35% B).

### Endocannabinoid analysis

For endocannabinoid analysis, samples were prepared in the same way as for Untargeted *N*-Acyl amide analysis. The chromatographic separation was performed on a Sciex exion (AB Sciex Inc., Framingham, MA) consisting of a binary pump, an autosampler and a thermostated column compartment. The column used was an XBridge BEH C18 2.5µm, 2.1×150mm column with a pre-column made with the same material. The eluents were A: 0.1 % FA and 1% ammonium acetate (1M) in water and B: 0.1 % FA and 1 % ammonium acetate (1M) in ACN/IPA (50:50). The gradient is presented in Supplementary Table 2, the injection volume was 1 µL, the flow rate was 0.4 mL/min and the column oven temperature was 40 °C. The detection was performed on a Sciex 7500 QTrap operating in MRM mode. The parameters used are presented in Supplementary Table 1 and 3. Quantification was performed using calibration curves from 0.01 ppb to 80 ppb (0.1 and 800 ppb for Arachidonic acid (AA)) using the internal standard method. Quantification was performed with Sciex OS analytics.

### Data analysis

#### MZmine data processing

The LC-MS/MS data underwent feature extraction with MZmine 4 as described here ^31^, employing stringent signal-to-noise thresholds (Noice level 100) for both MS1 and MS2 levels. These features, characterized by their unique isotope patterns and MS2 spectra, were exported in. MGF and .CSV formats for further analysis. All batch files and corresponding output files produced by processing the example datasets are available in the Supplementary data file.

#### Feature-Based Molecular Networking Workflow Description

The output files (. MGF and .CSV) generated from MZmine 4.4.3 were utilized in the Feature-Based Molecular Networking workflow on the GNPS2 platform (https://gnps2.org/homepage). For the analysis, the precursor ion mass (MS1) and MS2 fragment ion tolerances were set at 0.005 Da each. Connections between nodes were established when MS/MS spectral comparisons showed a cosine similarity of 0.6 or higher, with at least four matching fragment ions required to form the molecular network. All spectra in the networks were matched against the GNPS spectral libraries, applying a cosine threshold of 0.6 and a minimum of four MS2 matches. The processing task is accessible at ([Link to the GNPS task](https://gnps2.org/status?task=84896d95a4e149f9974ca8c3cf3e0905)) Additionally, molecular networks were visualized in Cytoscape software.

#### Reanalysis of public dataset (MSV000094551)

The raw MS/MS spectra were converted to mzML files using MSconvert (ProteoWizard), followed by feature extraction using MZmine 4.2.0. The feature quantification table was subsequently filtered with R scripts to remove features with average peak areas in samples lower than 3-fold of those in blanks. Metabolite annotations were performed with the default GNPS Spectral Libraries (including the N-acyl lipid library) and GNPS-BILE-ACID-MODIFICATIONS library using the Feature-Based Molecular Networking (FBMN) workflow on GNPS2. The spectra were searched against the GNPS libraries with precursor and fragment ion mass tolerances of 0.02 Da and 0.05 Da, respectively, with a cosine score threshold of 0.6 and a minimum of 4 matched peaks. The GNPS2 job is available at: https://gnps2.org/status?task=4e5f76ebc4c6481aba4461356f20bc35.

#### Statistical Analysis

The differences in the metabolite features between various vessels were analyzed using a multivariate linear model with the MaAsLin2 package in R (lipids ∼ Vessels), using media samples as the reference. Spearman’s rank correlation coefficients were calculated with the cor() function in R, specifying the method as ‘spearman’. The individual Spearman correlation coefficients, along with a bar plot and pie chart, were visualized as a heatmap using the ggplot2 package in the R statistical programming language. Fold change was calculated as the ratio of median peak areas in Intestin versus Feces samples with a pseudocount of 1 added to avoid division by zero. FC = (median_Intestin + 1) / (median_Feces + 1).

## Author Contributions

Conceptualization S.L.; methodology K.S., M.K., I.T., A.M.D, software, I.T., K.S., formal analysis and investigation, I.T., K.S., I.M., M.K., and S.L.; resources S.L.,A.M.D, M.O., T.H., P.W.P.G., V.C., H.M.R, P.D., S.D.F., and A.C.O; data curation, I.T., K.S., I.M.; writing—original draft preparation I.T., and S.L.; writing—review and editing, all authors.; visualization, S.L. and A.M.D; supervision; All authors have read and agreed to the published version of the manuscript.”

## Informed Consent Statement

The fecal samples used as inoculum in the colon simulator were given voluntarily by healthy adult Finnish volunteers. According to the Finnish law, no ethical approval was needed at the time the in vitro simulations were performed, since there was no interference with a person’s privacy, physical or mental integrity. “The study presented in the report is not considered medical research as defined in the Finnish Act on Medical Research (488/1999, as amended). Due to this, the in vitro study did not require an approval from the ethical committee and therefore such approval has not been obtained. In addition, as the study was not considered as medical research, the consent was not obtained in writing as required by the Act on Medical research, but orally.”

## Data Availability Statement

The metabolomics datasets generated in this study are available at MassIVE Repository (https://massive.ucsd.edu/ProteoSAFe/static/massive.jsp). MassIVE is a community resource developed by the NIH-funded Centre for Computational Mass Spectrometry. The data can be accessed directly at GNPS/MassIVE under the accession number MSV000097967.

## Acknowledgements

We thank the Turku Metabolomics Centre for the assistance and resources in the analyses of metabolites in this dataset. We would like to thank Tito Damiani for suggestions on the mzmine batch file. This work was supported by Research Council of Finland funding no. 363417 to S.L. M.O. was supported by Inflammation in human early life: targeting impacts on life-course health” (INITIALISE) consortium funded by the Horizon Europe Program of the European Union under Grant Agreement 101094099. A.M.D was supported by Research Council of Finland funding no. 347924. P.C.D was supported by NIH R01 DK136117 and H.M.R was supported by U24DK133658.

## References

1 Lamichhane, S., Sen, P., Dickens, A. M., Orešič, M. & Bertram, H. C. Gut metabolome meets microbiome: A methodological perspective to understand the relationship between host and microbe. Methods 149, 3–12, doi:10.1016/j.ymeth.2018.04.029 (2018).

2 Rooks, M. G. & Garrett, W. S. Gut microbiota, metabolites and host immunity. Nat Rev Immunol 16, 341–352, doi:10.1038/nri.2016.42 (2016).

3 Agus, A., Clément, K. & Sokol, H. Gut microbiota-derived metabolites as central regulators in metabolic disorders. Gut 70, 1174–1182, doi:10.1136/gutjnl-2020-323071 (2021).

4 Dekkers, K. F. et al. Author Correction: An online atlas of human plasma metabolite signatures of gut microbiome composition. Nat Commun 14, 2971, doi:10.1038/s41467-023-38607-1 (2023).

5 Nicholson, J. K. et al. Host-gut microbiota metabolic interactions. Science 336, 1262–1267, doi:10.1126/science.1223813 (2012).

6 Brown, E. M., Clardy, J. & Xavier, R. J. Gut microbiome lipid metabolism and its impact on host physiology. Cell Host Microbe 31, 173–186, doi:10.1016/j.chom.2023.01.009 (2023).

7 Lamichhane, S. et al. Linking Gut Microbiome and Lipid Metabolism: Moving beyond Associations. Metabolites 11, doi:10.3390/metabo11010055 (2021).

8 Gentry, E. C. et al. Reverse metabolomics for the discovery of chemical structures from humans. Nature 626, 419–426, doi:10.1038/s41586-023-06906-8 (2024).

9 Mannochio-Russo, H. et al. The microbiome diversifies long-to short-chain fatty acid-derived N-acyl lipids. Cell 188, 4154–4169.e4119, doi:10.1016/j.cell.2025.05.015 (2025).

10 Kenny, D. J. et al. Cholesterol Metabolism by Uncultured Human Gut Bacteria Influences Host Cholesterol Level. Cell Host Microbe 28, 245–257.e246, doi:10.1016/j.chom.2020.05.013 (2020).

11 Li, C. et al. Gut microbiome and metabolome profiling in Framingham heart study reveals cholesterol-metabolizing bacteria. Cell 187, 1834–1852.e1819, doi:10.1016/j.cell.2024.03.014 (2024).

12 Lee, M. T. et al. Gut bacterial sphingolipid production modulates dysregulated skin lipid homeostasis. bioRxiv, doi:10.1101/2024.12.29.629238 (2024).

13 Brown, E. M. et al. Bacteroides-Derived Sphingolipids Are Critical for Maintaining Intestinal Homeostasis and Symbiosis. Cell Host Microbe 25, 668–680.e667, doi:10.1016/j.chom.2019.04.002 (2019).

14 Mohanty, I. et al. The underappreciated diversity of bile acid modifications. Cell 187, 1801–1818.e1820, doi:10.1016/j.cell.2024.02.019 (2024).

15 Zierer, J. et al. The fecal metabolome as a functional readout of the gut microbiome. Nat Genet 50, 790–795, doi:10.1038/s41588-018-0135-7 (2018).

16 Kråkström, M. et al. Dynamics of the Lipidome in a Colon Simulator. Metabolites 13, doi:10.3390/metabo13030355 (2023).

17 Lamichhane, S. et al. Gut microbial activity as influenced by fiber digestion: dynamic metabolomics in an in vitro colon simulator. Metabolomics 12, 25, doi:10.1007/s11306-015-0936-y (2016).

18 Forssten, S. & Ouwehand, A. C. Dose-Response Recovery of Probiotic Strains in Simulated Gastro-Intestinal Passage. Microorganisms 8, doi:10.3390/microorganisms8010112 (2020).

19 Dührkop, K. et al. SIRIUS 4: a rapid tool for turning tandem mass spectra into metabolite structure information. Nature Methods 16, 299–302, doi:10.1038/s41592-019-0344-8 (2019).

20 Wang, M. et al. Sharing and community curation of mass spectrometry data with Global Natural Products Social Molecular Networking. Nature Biotechnology 34, 828–837, doi:10.1038/nbt.3597 (2016).

21 Charron-Lamoureux, V. et al. A guide to reverse metabolomics—a framework for big data discovery strategy. Nature Protocols, doi:10.1038/s41596-024-01136-2 (2025).

22 Duhrkop, K. et al. Systematic classification of unknown metabolites using high-resolution fragmentation mass spectra. Nat Biotechnol 39, 462–471, doi:10.1038/s41587-020-0740-8 (2021).

23 Dührkop, K. et al. Systematic classification of unknown metabolites using high-resolution fragmentation mass spectra. Nature Biotechnology 39, 462–471, doi:10.1038/s41587-020-0740-8 (2021).

24 Shamburek, R. D. & Schubert, M. L. Control of gastric acid secretion. Histamine H2-receptor antagonists and H+K(+)-ATPase inhibitors. Gastroenterol Clin North Am 21, 527–550 (1992).

25 Sandoval, M. & Shah, D. D. Diversity and prevalence of amino acid decarboxylase enzymes in the human gut microbiome – a bioinformatics investigation. bioRxiv, 2024.2012.2006.627230, doi:10.1101/2024.12.06.627230 (2024).

26 Tofalo, R., Cocchi, S. & Suzzi, G. Polyamines and Gut Microbiota. Front Nutr 6, 16, doi:10.3389/fnut.2019.00016 (2019).

27 Nam, Y. D., Kim, H. J., Seo, J. G., Kang, S. W. & Bae, J. W. Impact of pelvic radiotherapy on gut microbiota of gynecological cancer patients revealed by massive pyrosequencing. PLoS One 8, e82659, doi:10.1371/journal.pone.0082659 (2013).

28 Arul Prakash, S. & Kamlekar, R. K. Function and therapeutic potential of N-acyl amino acids. Chem Phys Lipids 239, 105114, doi:10.1016/j.chemphyslip.2021.105114 (2021).

29 Chesher, G. B. et al. The effect of cannabinoids on intestinal motility and their antinociceptive effect in mice. Br J Pharmacol 49, 588–594, doi:10.1111/j.1476-5381.1973.tb08534.x (1973).

30 Charron-Lamoureux, V. et al. A guide to reverse metabolomics-a framework for big data discovery strategy. Nat Protoc, doi:10.1038/s41596-024-01136-2 (2025).

31 El Abiead, Y. et al. Enabling pan-repository reanalysis for big data science of public metabolomics data. Nature Communications 16, 4838, doi:10.1038/s41467-025-60067-y (2025).

32 Tronel, A. et al. Pilot Study: Safety and Performance Validation of an Ingestible Medical Device for Collecting Small Intestinal Liquid in Healthy Volunteers. Methods and protocols 7, doi:10.3390/mps7010015 (2024).

33 Tronel, A. et al. Untargeted and semi-targeted metabolomics approach for profiling small intestinal and fecal metabolome using high-resolution mass spectrometry. Metabolomics 21, 84, doi:10.1007/s11306-025-02288-2 (2025).

34 Mannochio-Russo, H. et al. The microbiome diversifies long-to short-chain fatty acid-derived N-acyl lipids. Cell (2025).

